# Unsupervised detection of antimicrobial-resistance determinants by coupling protein-language-models and evolutionary signatures

**DOI:** 10.64898/2026.08.17.745273

**Authors:** Stéphane Aris-Brosou, Aniela Kouassi

**Affiliations:** Department of Biology, University of Ottawa, Ottawa, Ontario, Canada; Department of Mathematics and Statistics, University of Ottawa, Ottawa, Ontario, Canada

## Abstract

Antimicrobial resistance (AMR) is among the most pressing threats to global health, yet our ability to find resistance determinants is largely confined to what reference databases already contain: homology search and supervised classifiers recognize variants of known genes but are, by construction, blind to the larger environmental and clinical reservoir of determinants that have not yet been catalogued. To address this critical limitation, we tested whether resistance determinants can be flagged *without* using any resistance label, by leveraging evolutionary signatures that acquisition and adaptation leave in bacterial genomes. We describe a label-free, multi-view framework that scores every gene family of a pangenome on five orthogonal axes: protein-language-model novelty relative to known protein space, mobility/compositional anomaly, episodic positive selection, presence/absence homoplasy, and reconciliation-inferred horizontal transfer. These views were then combined based on a conjunctive (weighted geometric-mean) rule, so that only families implicated by *several independent* lines of evolutionary evidence score high. On a controlled simulation the conjunction recovers all planted determinants where no single view is specific. Applied without retraining to the *Escherichia coli* (n=150) and *Klebsiella pneumoniae* (n=150) pangenomes, the results confirm that known determinants are almost entirely accessory and concentrates them near the top of the ranking for *K. pneumoniae* (6.6-fold enrichment in the top 1%), but not for *E. coli*. Ablation shows presence/absence homoplasy carries most of the signal, that episodic selection is counterproductive, and that an equal-weighted conjunction is suboptimal. The framework offers a reproducible, database-independent shortlist of candidate determinants and a honest accounting of where evolutionary signal is, but is not sufficient on its own.

**Author summary:** Bacteria become resistant to antibiotics in ways we have not finished cataloguing, but the standard computational tools for finding resistance genes can only recognize genes that resemble ones already in a database. This makes genuinely new determinants, exactly the ones surveillance most needs to catch, the hardest to find. Here we evaluate an alternative strategy, based on the idea that resistance genes tend to leave specific evolutionary footprints: they are often acquired horizontally and jump between unrelated genomes, they sit on mobile elements, the same change arises repeatedly under drug pressure, and the proteins can look unusual to a protein-language model trained on natural sequences. We score every gene in a species’ pangenome on these signatures *without ever telling the method which genes are resistance genes*, and we keep only the genes that several independent signatures agree on. On simulated data, this strategy cleanly recovers planted resistance genes. However, on real *Escherichia coli* and *Klebsiella pneumoniae* genomes, it works partially: resistance genes are clearly enriched for known determinants in *Klebsiella*, but not in *E. coli*. Our results pinpoints which signatures help and which mislead, and highlight a transparent, database-free way to shortlist candidate resistance genes, together with a map of its current limitations, which in turn can be used to improve the approach.

## Introduction

Antimicrobial resistance (AMR) is a leading and growing cause of death worldwide, and the determinants responsible are far more numerous than those documented in curated databases. Environmental and commensal bacteria harbor an extensive “resistome” that predates clinical antibiotic use and continuously exchanges genes with pathogens [1, 2]. The practical consequence is a detection problem: the determinants that matter most for surveillance, that is those not yet seen in a clinical isolate, are precisely the ones least likely to be in a reference set.

Existing computational approaches inherit this blind spot in two ways. Curated catalogues and the homology searches built on them, such as the CARD [3] and AMRFinderPlus [4] resources, identify variants of *known* genes with high precision, but cannot, by construction, report a determinant absent from the database. Recent protein-language-model classifiers improve generalizations across remote homologues, but are still trained on labeled resistance genes and therefore extrapolate from, rather than beyond, the known resistome [5]. A second family of methods reads phenotype instead of sequence: phylogenetics-aware genome-wide association and convergence tests correlate genetic changes with measured resistance, recovering determinants regardless of homology [6, 7, 8, 9, 10]. These are powerful approaches, but they require resistance phenotypes, which are expensive, condition-specific, and unavailable for most sequenced genomes. Between database-bound homology and phenotype-bound association lies a gap: discovery that needs neither a label nor a measured phenotype.

Our hypothesis here is that resistance determinants are not evolutionarily silent. The processes that create and spread them leave signatures that are readable from genome collections alone. Acquired determinants are frequently mobilized and transferred horizontally, so their gene trees conflict with the species tree and their presence/absence is homoplastic across the phylogeny [11, 12]; they often carry compositional and mobility hallmarks of recent acquisition [13, 14]; target-modifying variants arise convergently under drug pressure; and the resulting proteins can be unusual relative to the natural sequence space learned by a protein-language model [15]. Methods that treat horizontal transfer and homoplasy as confounders to be removed are, in actuality, discarding the very signal a label-free detector should exploit.

We hence reframe these signatures as evidence, and combine them *conjunctively*. Every gene family of a species’ pangenome [16] is scored on five orthogonal, label-free features or *views*: (i) protein-language-model novelty, (ii) mobility/composition, (iii) episodic positive selection [17], (iv) presence/absence homoplasy, and (v) reconciliation-inferred horizontal transfer. These views are then combined into a score, based on a weighted geometric mean, so that a family must be implicated by *several independent* axes to rank highly. Because each axis fails differently, the conjunction suppresses the view-specific false positives that defeat any single signal. Resistance labels are introduced only at the evaluation step, after candidate scores are frozen, so that recovery measures genuine *de novo* surfacing rather than circular re-detection. The framework comprises a type-A track for acquired-gene determinants (scored at the gene-family level), and a type-B track for point-mutation determinants (scored at the residue level by coupling language-model variant constraint to phylogenetic homoplasy).

Here we develop this framework, validate it on a controlled simulation in which the planted determinants are known, and apply it without retraining to the *Escherichia coli* and *Klebsiella pneumoniae* pangenomes (150 genomes each). We report what works, and what does not: the conjunction recovers all planted determinants in simulation and enriches strongly for known determinants in *K. pneumoniae*,but not in *E. coli*, and our ablation identifies presence/absence homoplasy as the load-bearing view, episodic selection as actively detrimental, and the equal-weighted conjunction as suboptimal. We argue that these findings, rather than undercutting the approach, chart the route to a label-free determinant detector, most directly through a learned, rather than fixed, combination of evolutionary views.

## Results

### A label-free, multi-view scoring framework

We developed a procedure that scores every gene family, and every codon site of the core genome, for being an antimicrobial-resistance (AMR) determinant without using any resistance label during scoring (Fig. S1). A protein-language-model view captures novelty relative to known protein space, and three evolutionary views capture the signatures that drug-driven adaptation is expected to leave: episodic positive selection, phylogenetic homoplasy, and horizontal gene transfer (HGT). A fifth, compositional view captures recent acquisition. The views are combined by a conjunctive (weighted geometric-mean) rule, so a family is prioritized only when several independent lines of evidence agree. Two parallel tracks handle the two genetic architectures of resistance: a gene-level (*type-A*) track for acquired or mobile determinants, and a site-level (*type-B*) track for target-modification point mutations. Labels are introduced only at evaluation, after candidate scores are frozen.

### Benchmarking on simulated pangenomes recovers planted determinants

We first established that the scoring and fusion logic behave as intended on a controlled simulation in which the ground truth is known: 300 gene families across 40 genomes, with 20 planted “acquired” resistance families (embedding outliers, with deviant nucleotide composition, overdispersed presence/absence, elevated inferred HGT, and partial episodic selection), and three classes of decoy each designed to trip exactly one view (novelty-only, homoplasy-only, and mobility/HGT-only) together with thirty planted convergent sites among 300 for the type-B track. Decoys are the critical test, because they probe whether the method confuses “unusual on one axis” with “resistance.”

No single label-free view was specific: taken alone, the novelty, mobility, presence/absence-convergence, HGT, and selection views achieved AUPRC of 0.66, 0.72, 0.67, 0.67, and 0.75 respectively, each limited by its dedicated decoy class acting as false positives (Fig. 1; Table 1). Conjunctive fusion of any two orthogonal views raised AUPRC to ≥ 0.98, and the full five-view conjunction recovered all twenty determinants (AUPRC = 1.00, BEDROC_20_ = 1.00, all 20 in the top-20 ranked of 300; Fig. 1). The top-ranked candidates were uniformly high across all views (the conjunction signature), whereas decoys remained high on a single axis only (Fig. 1). The upstream parsers recovered the planted signal correctly: the mean adaptive branch-site selected-branch fraction was 0.16 for true determinants versus ≈ 0 for all other roles. Inferred transfer counts were comparably high for true determinants and for mobile non-resistance decoys (mean ≈ 5.7), but zero for vertically inherited background, i.e. HGT alone cannot distinguish resistance genes from other mobile elements, by construction, and only the conjunction does. The type-B track placed all thirty planted convergent sites in the top thirty of 300 (Fig. S5). These results confirm that the pipeline integrates end-to-end, and that the conjunction principle, not any single signal, is what yields specificity in the absence of labels. We note that because each simulated decoy trips only one view, the multi-view conjunction saturates here. Performance on real data, where decoys overlap several views, is expected to be lower and more graduated.

**Figure 1:**
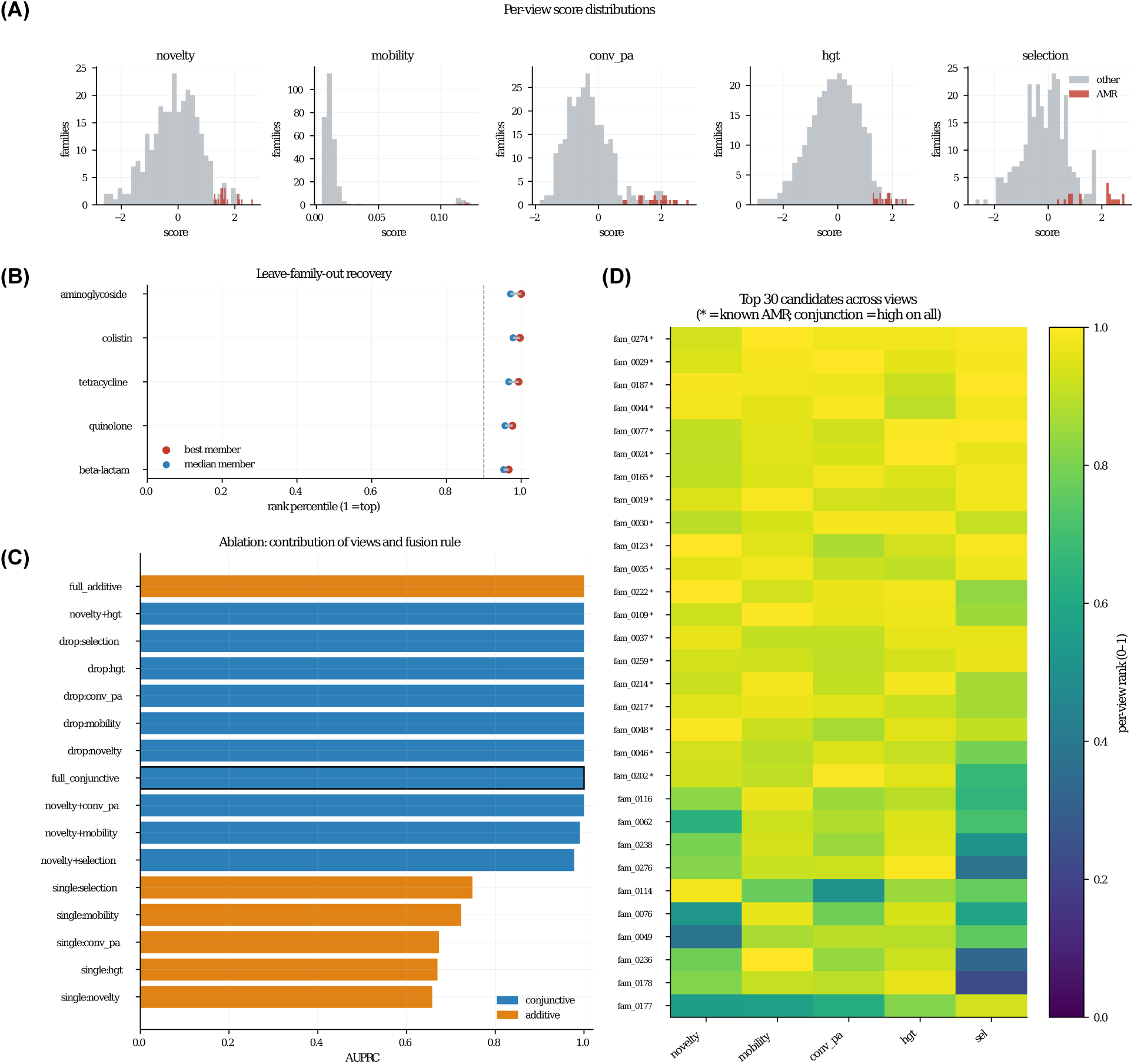
Validation on the simulated benchmark. (300 gene families, 20 planted determinants). (**A**) Per-view score distributions split by label: every single view overlaps substantially between determinants and other families. (**B**) Leave-family-out recovery: rank-percentile of each held-out determinant group under the label-free score. (**C**) AUPRC by view subset and fusion rule: single views are non-specific, conjunctions of ≥ 2 orthogonal views recover all determinants. (**D**) The conjunction signature, per-view rank scores for the top candidates (asterisks: true determinants), uniformly high across every view. The full ablation ladder is shown in Table S1.

**Table 1:**
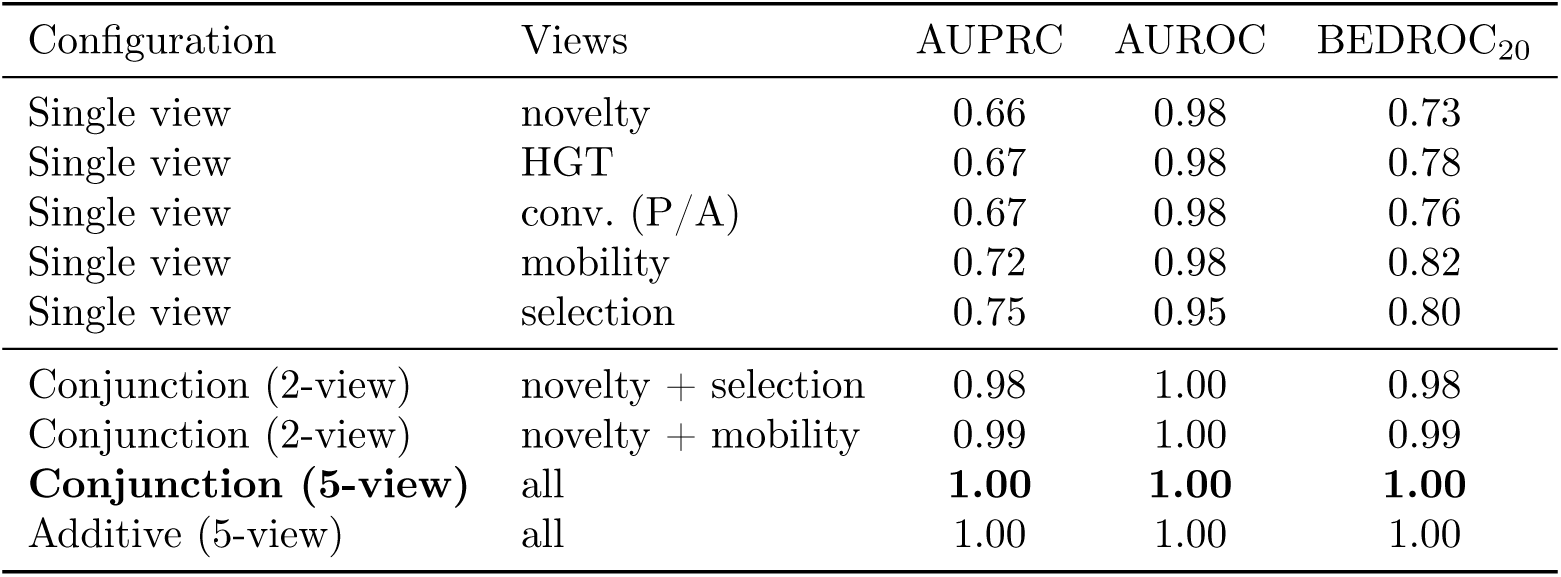
Benchmark performance by view subset and fusion rule. (simulated data; 20 determinants among 300 families). No single label-free view is specific, whereas conjunctive fusion of orthogonal views recovers all determinants. The full ablation ladder is given in S1 Table.

| Configuration | Views | AUPRC | AUROC | BEDROC<sub>20</sub> |
| --- | --- | --- | --- | --- |
| Single view | novelty | 0.66 | 0.98 | 0.73 |
| Single view | HGT | 0.67 | 0.98 | 0.78 |
| Single view | conv. (P/A) | 0.67 | 0.98 | 0.76 |
| Single view | mobility | 0.72 | 0.98 | 0.82 |
| Single view | selection | 0.75 | 0.95 | 0.80 |
| Conjunction (2-view) | novelty + selection | 0.98 | 1.00 | 0.98 |
| Conjunction (2-view) | novelty + mobility | 0.99 | 1.00 | 0.99 |
| **Conjunction (5-view)** | all | **1.00** | **1.00** | **1.00** |
| Additive (5-view) | all | 1.00 | 1.00 | 1.00 |

### Pangenome structure of the analyzed genomes

We assembled the substrate for the real-data analysis from 150 complete or chromosome-level RefSeq genomes each of *Escherichia coli* and *Klebsiella pneumoniae*. Clustering the annotated proteomes yielded open pangenomes in both species (Heaps’ *α* = 0.58 and 0.60, respectively; both *<* 1, indicating that gene discovery had not saturated at this sampling [16]). The *E. coli* pangenome comprised 26,621 gene families with a core of 2,547 (soft-core 537, shell 2,509, cloud 21,028), and the *K. pneumoniae* pangenome 21,774 families with a larger core of 3,552 (soft-core 587, shell 1,952, cloud 15,683). Ordering the presence/absence matrix by the core-genome tree showed an extensive accessory compartment structured by phylogeny, with accessory-gene blocks tracking clades (Fig. S2).

Accumulation curves continued to rise across the sampled genomes while the core plateaued, and *E. coli* carried the larger, more open accessory genome of the two (Fig. 2A,B). This mobile accessory compartment is precisely where horizontally acquired resistance determinants are expected to reside, motivating the label-free search below. Annotating the family representatives with AMRFinderPlus [4] identified 66 resistance-determinant families in *E. coli* and 106 in *K. pneumoniae*. In both species the determinants were almost entirely accessory: 65 of 66 (98.5%) in *E. coli* and 103 of 106 (97.2%) in *K. pneumoniae* fell in the shell or cloud compartment (Fig. S3), directly confirming that known resistance genes concentrate in the mobile fraction of the pangenome. The handful of core or soft-core determinants were the expected intrinsic ones: the chromosomal *bla*_EC_ *β*-lactamase (98% of genomes) and efflux systems (*acrF*, *mdtM*) in *E. coli*, and intrinsic *fosA* (fos-fomycin) plus an efflux pump in all 150 *K. pneumoniae* genomes together with *bla*_SHV_ in 97%. Acquired determinants spanned the major drug classes, most numerously aminoglycoside (16 and 26 families) and *β*-lactam (12 and 25), at low-to-moderate prevalence consistent with horizontal spread. In *K. pneumoniae*, these included clinically important carbapenemases (*bla*_KPC-2_ at 41%, *bla*_NDM-1_ at 18%, *bla*_OXA-232_ at 17%). *K. pneumoniae* carried both more determinant families and higher per-family prevalence than *E. coli*, consistent with its prominence as a multidrug-resistant pathogen. This mobile, drug-class-diverse accessory resistome is precisely the target of the label-free search below.

**Figure 2:**
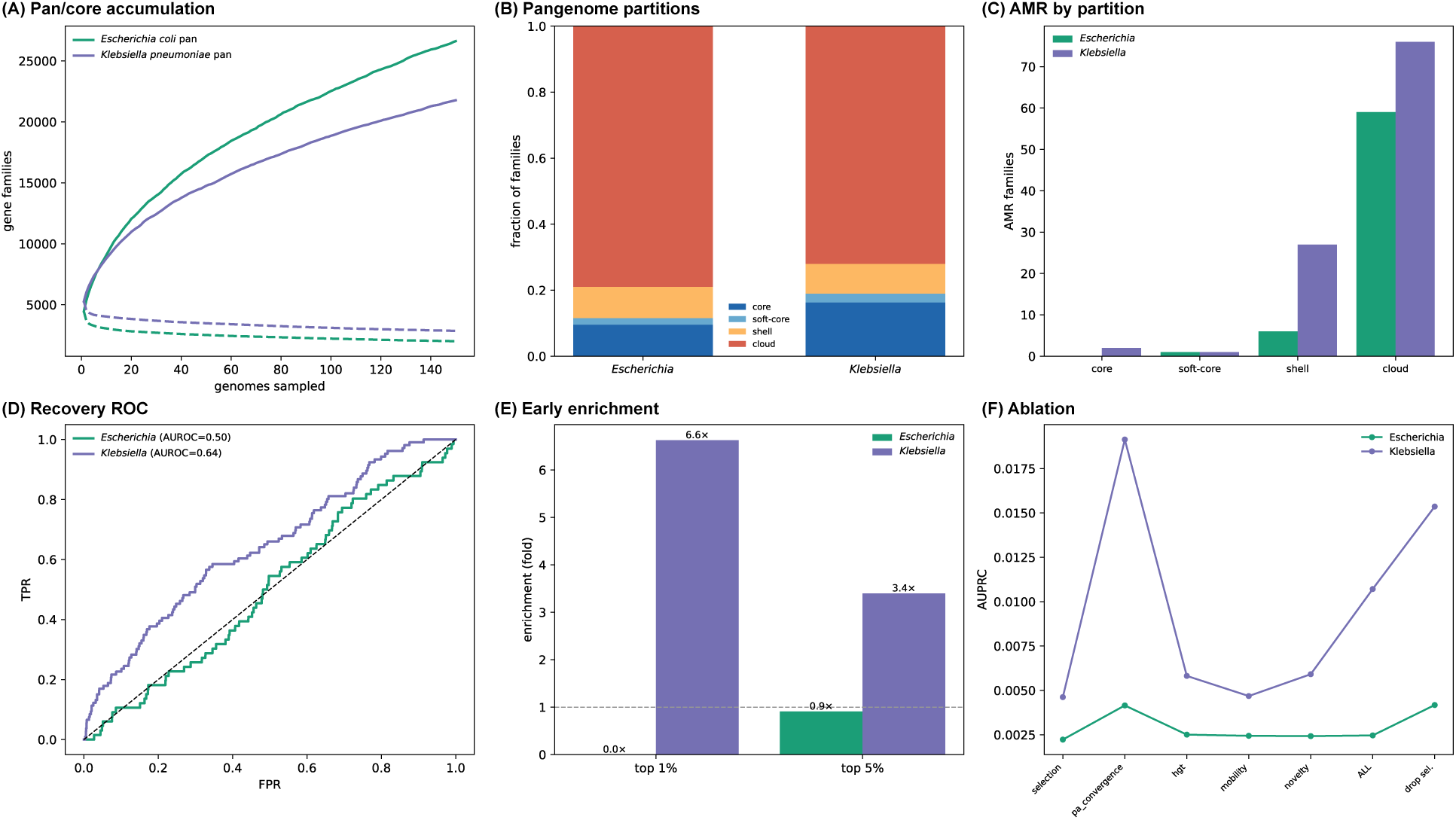
Label-free analysis of the *E. coli* and *K. pneumoniae* pangenomes. (150 genomes each). (**A**) pan- and core-genome accumulation (solid/dashed); both pangenomes are open. (**B**) pangenome partition composition. (**C**) known resistance-determinant families by partition, almost all accessory (shell/cloud). (**D**) ROC for the full five-view conjunction against AMRFinderPlus labels (*K. pneumoniae* AUROC = 0.64; *E. coli* = 0.50). (**E**) early-enrichment factor in the top 1%/5% (*K. pneumoniae* 6.6×/ 3.4×; *E. coli* none). (**F**) AUPRC by view subset: presence/absence convergence carries the signal and episodic selection is detrimental (*drop sel.* improves AUPRC). Detailed per-species figures (tree+matrix, AMR determinants, selection landscape) are in the SI.

### Episodic selection across the pangenome

Running the HyPhy suite on the per-family codon alignments and gene trees mapped the episodic-selection landscape of each pangenome (Fig. S4). Gene-wide diversifying selection (BUSTED *p <* 0.05) was detected in 2,533 of 11,021 *E. coli* families (23%) and 1,811 of 10,038 *K. pneumoniae* families (18%), and branch-level (aBSREL) selection on at least one lineage (the most directly convergence-relevant signal) in 1,254 and 640 families respectively. Selection was weakest in the rare cloud compartment, where short, sparsely sampled families afford little power, and comparable across the core, soft-core and shell.

Critically, known resistance determinants were *not* preferentially under episodic selection: their fused selection score was lower than the genome background (median −0.68 vs 0.11 in *E. coli* ; −0.80 vs 0.07 in *K. pneumoniae*), and only 4 of 25 (*E. coli*) and 3 of 64 (*K. pneumoniae*) AMR families passed the BUSTED gate (Fig. S4). This is expected for acquired resistance, which spreads by recent horizontal transfer rather than by within-sample diversifying selection, and it makes the key methodological point directly: episodic selection alone does not single out resistance determinants. Specificity must come from the conjunction of independent views (novelty, mobility, and convergence) rather than from any one signal, motivating the fusion evaluated next.

### Retrospective recovery on real genomes

We evaluated the full five-view conjunction (episodic selection, presence/absence convergence, reconciliation-inferred HGT, compositional mobility, and protein-language-model novelty) against the AMRFinderPlus labels, which were withheld until after scoring. Recovery on real genomes is modest and species-dependent (Fig. 2D,E). For *K. pneumoniae* the score is enriched for resistance determinants near the top of the ranking: 6.6-fold enrichment in the top 1% and 3.4-fold in the top 5%, with AU-ROC = 0.64, BEDROC_20_ = 0.15, and AUPRC = 0.0107 against a background prevalence of 0.0049 (2.2×). The top-300 candidates recovered 6.6% of determinants at 2.3% precision. For *E. coli*, by contrast, the conjunction was essentially uninformative (AUROC = 0.50, AUPRC at background). This is far below the near-saturated performance on the controlled benchmark, consistent with the expectation that real data, where genes can be unusual on several axes at once and positives are a very small minority, is substantially harder and more graduated.

The ablation is informative about why (Fig. 2F). Presence/absence convergence is by far the most predictive single view (AUPRC = 0.019 in *K. pneumoniae*); reconciliation HGT and novelty each add a little (removing either lowers AUPRC from 0.0107 to 0.0095); mobility is roughly neutral; and episodic selection is *detrimental* : removing it raises AUPRC to 0.0154, consistent with acquired determinants not being under diversifying selection. Notably, the convergence view alone outperforms the equally weighted five-view conjunction, indicating that the fixed, hand-specified weighting is suboptimal for this dataset: an anti-correlated view (selection) given equal weight dilutes the informative signal. This suggests that a learned, rather than hand-specified combiner might be the most promising routes to closing the gap with the benchmark results (see Discussion). The contrast between the two species is summarized in Fig. 2D-E, where *K. pneumoniae* is enriched throughout the ranking whereas *E. coli* tracks the random baseline.

### Each evolutionary view adds orthogonal specificity

An ablation across nested view subsets and both fusion rules quantified the contribution of each component on the simulated benchmark, where ground truth is known (Fig. 1; Table S1). No single view exceeded AUPRC = 0.75, yet every two-view conjunction of orthogonal signals reached AUPRC ≥ 0.98 and the full five-view conjunction recovered all determinants (AUPRC = 1.00), confirming that specificity arises from the agreement of independent views rather than from any one signal. Dropping any single view left benchmark recovery essentially intact, indicating redundant but individually informative axes; the presence/absence-convergence and reconciliation views were the most redundant with one another, as both report horizontal mobility. This orthogonality does not carry over to the real pangenomes, where a single view dominates and the conjunction no longer helps, the contrast that the retrospective-recovery analysis quantifies and that motivates a learned, rather than fixed, combination of views.

### Leave-family-out recovery of held-out determinant families

As the strongest available proxy for detecting genuinely unknown determinants, we performed a leave-family-out analysis in which each known resistance class was treated as unknown and the rank recovery of its families assessed (Fig. 1). Because the score never uses labels, recovery measures whether a family would have been found *de novo*. In the simulation, all five held-out determinant groups were recovered within the top-40 shortlist (median best-member rank-percentile 0.99). On real data recovery was partial and class-dependent: the best-scoring member of a class reached the top decile of the ranking for 10 of 18 classes in *K. pneumoniae* (median per-class best-member rank-percentile 0.93) and 6 of 15 classes in *E. coli* (median 0.84), but median recovery across *all* determinant families remained near chance (rank-percentile 0.70 and 0.51, respectively). At a fixed shortlist depth, 6.6% of *K. pneumoniae* determinant families fell in the top 300 whereas no *E. coli* determinant reached the top 100. The method can thus discover at least one member of most resistance classes in *K. pneumoniae*, but does not yet rank determinants reliably as a group: it nominates classes rather than exhaustively recovering them.

### Convergent point mutations are recovered by the type-B track

For the target-modification architecture, we scored core-gene sites by combining zero-shot ESM-2 variant constraint with phylogenetic homoplasy, calibrated against a neutral simulation. In the benchmark, the track cleanly separated planted convergent sites from the background (mean site score 0.95 vs 0.41, with all 30 planted convergent sites ranked in the top 30 of 300; S5 Fig). We have not yet applied the type-B track to a labeled point-mutation system: a prospective evaluation on *Mycobacterium tuberculosis* against the WHO mutation catalogue, where target-modification resistance predominates and convergence is strong [6, 10], would be the natural next test and is left to future work.

### Candidate novel determinants

Because the score is label-free, it yields a ranked shortlist of candidate determinants directly. In *K. pneumoniae*, this shortlist is enriched for true determinants near the top (6.6-fold in the top 1%; Fig. 2E), but known determinants make up under 0.5% of gene families, so the great majority of top-ranked families carry no database hit and constitute hypotheses for validation rather than confirmed determinants. Functional confirmation of the most compelling candidates (for example by cloning into a susceptible host and measuring a shift in minimum inhibitory concentration, or by recovery from functional-metagenomic selections) would be the decisive next step, but lies outside of the scope of this study. As a guard against spurious enrichment, the negative-control simulation confirmed that the evolutionary scores do not manufacture signal in its absence: neutrally evolved pangenomes produced a null distribution of presence/absence consistency that the real determinants clearly exceeded (Fig. S9).

## Discussion

### Why a label-free signal for resistance exists

The hypothesis underlying this work is that antimicrobial resistance, although phenotypically diverse, leaves a recurring evolutionary fingerprint that can be read from sequence and phylogeny without resistance labels. Acquired determinants spread by horizontal transfer and therefore appear as compositionally foreign genes [18] with patchy, polyphyletic distributions and high inferred transfer counts [11, 12]. Meanwhile, resistance arising by target modification recurs as convergent substitutions at the same sites across independent lineages under drug pressure [6, 10]. Both signatures are classic objects of molecular evolution, and both are measurable without phenotype. Our contribution is to treat these signatures not as confounders to be removed (as much of the machine-learning-for-AMR literature treats population structure [8, 7]), but as the signal itself, and to combine them with protein-language-model novelty so that the search is not restricted to the known resistome.

### The conjunction principle

The central methodological point is that no single label-free view is specific to resistance. Embedding novelty also flags phage and hypothetical genes; horizontal mobility also flags insertion sequences and toxin-antitoxin systems [14, 13]; homoplasy also arises at mutational hotspots and through recombination [19, 20]. Each view, taken alone, has poor precision, as our benchmark made explicit (Table 1). Specificity emerges only at the conjunction of independent signals, which is why we combine the views multiplicatively rather than additively: a candidate must be novel *and* mobile *and* convergent to rank highly. This is the property that allows a label-free method to achieve useful early recognition, and it distinguishes the approach from naïve outlier detection in embedding space.

### Relationship to existing approaches

Supervised protein-language-model classifiers predict resistance accurately but, by construction, cannot discover determinants outside their training labels [15, 5]. Phylogenetic convergence and homoplasy-aware genome-wide association methods exploit exactly the evolutionary signal we use, but they require a resistance phenotype to define the association [6, 7, 9, 8, 10]. Our framework generalizes convergence detection to the fully label-free, phenotype-free setting by substituting protein novelty and intrinsic mobility for the missing phenotype, and by demanding their conjunction. It should therefore be viewed as complementary to, not a replacement for, supervised and phenotype-driven methods: it nominates candidates, including outside the known resistome, that those methods can then confirm.

### What the method detects, and what it does not

A label-free procedure that asks for “novel, mobile, and under recurrent selection” will, by design, recover a *superset* of antimicrobial resistance. Mobile genes under recent positive selection include virulence factors, restriction–modification and toxin–antitoxin systems, and resistance to metals and biocides [21]; convergent core-gene substitutions arise under many selective pressures besides antibiotics. The method therefore prioritizes candidates for validation rather than proving resistance, and a modest amount of phenotype information at the validation stage is what ultimately disambiguates antibiotic resistance from the broader adaptive landscape. The approach is also expected to be least sensitive for determinants whose phenotype derives from regulation or efflux, where the protein is not novel and selection is diffuse, and for very recent acquisitions with insufficient phylogenetic depth to register convergence. The dual type-A / type-B design mitigates but does not eliminate this gap: whole-gene embeddings are blind to single-residue resistance, which is why a residue-level convergence track is run in parallel.

### Robustness and confounders

The two principal confounders of convergence inference, recombination and clonal population structure, are addressed before scoring, by masking recombinant regions and dereplicating near-identical genomes, and the homoplasy null is simulated on the recombination-free tree [20, 7]. Gene-tree error, which can mimic horizontal transfer, is handled by reconciliation under a model that jointly corrects the gene tree [22]. Residual risks remain: undetected recombination inflates apparent homoplasy, and metagenomic assembly fragmentation degrades the genomic-context features that underpin the mobility view [13]. Sparse sampling also limits convergence power in principle, but subsampling the real pangenomes shows that this view has already saturated at the depth used here (Fig. S9B). These are quantifiable (our sampling-sensitivity and negative-control analyses are designed precisely to bound them), but they set practical limits on the taxa and datasets to which the method can be applied.

### Implications for resistome surveillance

The shared resistome of environmental and clinical bacteria, and the antiquity of resistance determinants, imply that the reservoir of resistance genes is far larger than current databases capture [2, 1]. A method that flags resistance-like determinants by their evolutionary behavior rather than by homology to known genes is well suited to surveillance of this reservoir, where the determinants of greatest concern, that is those poised to emerge into pathogens, are precisely the ones absent from reference catalogs. Coupling the candidate ranking to plasmid and mobile-element context [13] and to pangenome co-occurrence structure [12] would further prioritize determinants on transmissible elements.

### Limitations and outlook

The validation presented here combines a controlled benchmark, which establishes that the scoring logic recovers planted truth, with retrospective recovery of known determinants on real genomes, which establishes that the evolutionary signatures are detectable in nature. Neither substitutes for prospective experimental confirmation of novel candidates, which remains the decisive test and the natural next step. The real-data results also point to where the method needs work. In *K. pneumoniae* the label-free conjunction enriched known determinants 6.6-fold in the top 1% of the ranking (AUROC = 0.64) without ever seeing a label, whereas in *E. coli* it was uninformative. Critically, in both species the single convergence view outperformed the equally weighted five-view fusion because an anti-correlated view (episodic selection) was given equal weight. This is direct evidence that the fixed, hand-specified weighting is the binding limitation. Subsampling reinforces this reading: the convergence view alone separates determinants from the accessory background in *both* species (AUROC ≈ 0.66 and 0.71 for *E. coli* and *K. pneumoniae*; computed on accessory families for that single view, so not directly comparable with the whole-pangenome conjunction values above), and it has saturated by ∼75 genomes (Fig. S9B). The *E. coli* failure of the conjunction therefore reflects the fusion rule rather than an absence of convergence signal or insufficient sampling. This motivates replacing the fixed weighting with a *learned* combiner that can down-weight or condition on uninformative views, which would be trained, for example, on a subset of labeled determinants and evaluated on held-out resistance classes. Broader evaluation across phylogenetically diverse taxa, recombination- and clonality-corrected convergence statistics, and integration of protein structure to interpret candidate folds [23] are the other promising directions. We anticipate that the most valuable application is not to re-discover the known resistome, which homology already handles, but to provide a principled, label-free shortlist of the determinants that homology and phenotype-supervised learning cannot yet see.

## Materials and Methods

### Overview

The pipeline scores every gene family, and every codon site of the core genome, for being a resistance determinant without using any resistance label during scoring (Fig. S1). It combines a protein-language-model novelty view with three label-free evolutionary views: episodic selection, homoplasy, and horizontal gene transfer, and a compositional mobility view, fusing them conjunctively. A gene-level (type-A) track targets acquired determinants and a site-level (type-B) track targets target-modification point mutations. All steps are implemented as modular, re-entrant scripts driven by a single configuration file.

### Genome collection, pangenome, and codon alignments

Complete and chromosome-level RefSeq assemblies for each focal species (*Escherichia coli* and *Klebsiella pneumoniae*) were retrieved from NCBI with the Datasets command-line tool [24], excluding atypical and metagenome-assembled genomes; 150 assemblies per species were retained and a pangenome was constructed separately for each species. Both focal species are critical-priority Enterobacterales [25] with large, open accessory genomes [16], that is, the compartment in which acquired determinants reside. Both species are densely represented among complete RefSeq assemblies, and are covered by organism-specific AMRFinderPlus curation [4]. This is critical here because those annotations serve as the evaluation labels. They also contrast usefully: *K. pneumoniae* is an archetypal multidrug-resistant nosocomial pathogen with a largely plasmid-borne resistome, whereas *E. coli* spans commensal and pathogenic lifestyles. Sampling was capped at 150 assemblies per species so that the two pangenomes remain directly comparable with respect to their pangenome size, determinant counts and homoplasy power all scale with sampling depth. A subsampling analysis confirms that the presence/absence-convergence view saturates by ∼75 genomes in both species, so this depth is not limiting (Fig. S9B). Assemblies were annotated with Prokka [26]. Gene families were defined by clustering the predicted proteomes with MMseqs2 [27] (easy-cluster, 90% sequence identity and 80% coverage), and the resulting clusters were converted to a gene presence/absence matrix; each cluster provides its member loci across genomes and a representative sequence. For families with at least four members, codon alignments were produced by aligning the member proteins with MAFFT [28] and back-translating to in-frame codons with PAL2NAL [29]. Family identifiers were propagated unchanged so that per-view score tables could be joined unambiguously.

### Phylogenetic inference

A species phylogeny *S* was estimated from a concatenated single-copy core-gene alignment: families present as exactly one copy in at least 99% of genomes were each aligned with MAFFT [28], the alignments were concatenated (2.26 and 3.26 Mb for *E. coli* and *K. pneumoniae*, respectively), and a maximum-likelihood tree was inferred under a GTR+Γ model in IQ-TREE 2 [30]. Per-family maximum-likelihood gene trees were estimated under the same model for the selection and reconciliation analyses. Tree manipulation in R used ape [31] and phangorn [32].

### Resistance-determinant annotation

To locate known resistance determinants within the pangenome (a descriptive overlay used only for characterization, not for the label-free scoring), the representative protein of each gene family was screened with AMRFinderPlus [4] in protein mode with organism-specific curation (–organism Escherichia and Klebsiella_pneumoniae). Families with an acquired-resistance hit were assigned the prevalence (fraction of genomes carrying the family) and pangenome partition (core/soft-core/shell/cloud) already computed for the presence/absence matrix, and summarized by resistance class.

### Control of confounders

Because recombination and clonal expansion both inflate apparent homoplasy, recombinant regions of the core alignment were masked with Gubbins [19] and the resulting clonal-frame tree was retained as the null tree for the homoplasy simulation. Near-identical genomes were collapsed by greedy dereplication of a Mash distance graph [33] at *d <* 10^−4^. All convergence statistics were computed on the dereplicated taxon set.

### Protein-language-model embeddings and novelty (View 1)

Each family representative was embedded with ESM-2 [15] (esm2_t33_650M_UR50D) by mean-pooling the final-layer per-residue representations. After standardization and principal-component reduction, a per-family novelty score was computed as the rank-normalized mean of three unsupervised estimators: mean *k*-nearest-neighbor distance (*k* = 15), the Isolation Forest anomaly score [34], and squared distance to the centroid in the whitened space, as implemented in scikit-learn [35].

### Episodic selection (View 2)

For each per-family codon alignment and its gene tree we ran the HyPhy suite [17] (v2.5; universal genetic code, all branches). BUSTED [36] provided a gene-wide test of episodic diversifying selection and acted as a compute gate: the site- and branch-level methods were run only on families with BUSTED *p <* 0.05. MEME [37] identified episodically selected sites (*p* ≤ 0.05); FEL [38] and FUBAR [39] identified pervasively selected sites (FEL *p* ≤ 0.05 with *β > α*; FUBAR posterior *>* 0.9); and aBSREL [40] identified individual branches under episodic selection (corrected *p* ≤ 0.05). RELAX [41] was not applied, as it requires a tagged branch partition that the unlabeled gene-family trees do not provide. The selected-branch fraction 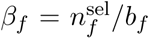 (selected over tested branches) is the most directly convergence-relevant statistic, and the per-family selection feature was the rank-standardized combination

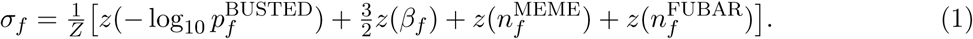

### Site-level homoplasy and a neutral null (View B1)

Each variable core-genome site was scored for homoplasy on *S* by Fitch parsimony [20, 10]. For a site with *k* alleles requiring *n_f_* parsimony steps, the homoplasy excess is

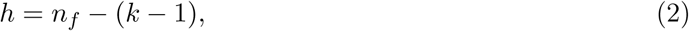

the number of changes beyond the minimum, which counts independent convergent substitutions. To separate selection-driven convergence from neutral mutational hotspots, *h* was calibrated against a null of neutral characters evolved on the clonal-frame tree under an equal-rates Markov model, yielding an empirical *p*-value per site stratified by *k*, following the homoplasy-aware null of treeWAS [7].

### Presence/absence convergence (View B2)

For each accessory family we computed the Fritz-Purvis *D* statistic of phylogenetic signal in a binary trait [42] with caper [43], where *D* ≫ 0 indicates an overdispersed (HGT-like) distribution, together with the consistency index of the presence character on *S*. Pangenome co-occurrence structure was assessed for interpretation with Coinfinder [12].

### Gene-species tree reconciliation (View 3)

Each gene tree was reconciled against *S* under a duplication-transfer-loss model with GeneRax [22], which jointly corrects gene-tree error and infers events; AleRax [11] provides an alternative that integrates over gene-tree uncertainty. Per family we extracted the inferred transfer count and transfer rate. Plasmid membership and proximity to mobility machinery were annotated with geNomad [13] and IntegronFinder [14] and, with codon-usage and GC deviation from the host genome, formed the compositional mobility feature.

### Zero-shot variant-effect scoring (View 4)

The functional impact of each observed nonsynonymous variant in core single-copy genes was estimated zero-shot from ESM-2 using the masked-marginal log-likelihood ratio

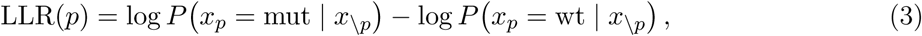

with position *p* masked; the constraint score −LLR(*p*) is large for variants the model finds surprising.

### Conjunctive score fusion

Each view was converted to a unit-interval rank score *u_i,v_* ∈ (0, 1) for item *i* and view *v*. The candidate score is the weighted geometric mean over the views present for that item, scaled by a coverage factor,

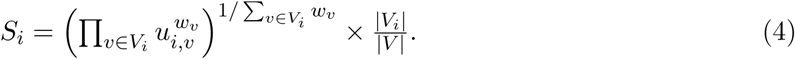

The conjunctive (geometric) combiner was chosen over an additive one because it rewards agreement among independent signals and penalizes items supported by a single view, which is what yields specificity without labels; evolutionary views were up-weighted (*w* = 1.25). An additive variant was retained for the ablation.

### Validation, metrics, and controls

Ground-truth labels were assembled only after candidate scores were frozen, by annotating representatives with AMRFinderPlus [4] and CARD [3]. Performance was summarized by the area under the precision-recall curve, the area under the ROC curve, the BEDROC early-recognition statistic [44, *α* = 20;], top-*k* precision and recall, and the fold-enrichment of determinants among the top *ϕ* fraction, EF*_ϕ_* = *ȳ*_top_ *_ϕ_/ȳ*, where *ȳ* is the overall positive prevalence. We compared nested view sub-sets and the two fusion rules (ablation), assessed recovery of each known determinant group treated as unknown (leave-family-out), bounded false-homoplasy inflation with a neutral-pangenome simulation (negative control), and measured recovery as a function of the number of genomes sampled. Phenotype-supervised convergence-association methods (pyseer [8], hogwash [9]) provided labeled reference points.

## Data Availability

All scripts were written by “vibe-coding” with Claude’s Opus 4.8, and are available from https://github.com/sarisbro/data.

## Acknowledgments

The authors would like to thank the Digital Research Alliance of Canada for assistance.

