## Supplementary material for "Unsupervised detection of antimicrobial-resistance determinants by coupling protein-language-models and evolutionary signatures": SI file

### Supplementary methods and notes

#### Conjunctive fusion score

Each label-free view $v$ is converted to a rank-normalized score $u_v \in (0, 1)$ (1 = most determinant-like), so that the views are comparable and robust to their differing native scales. For a gene family with the set of available views $V$ (some views may be missing for a given family), the conjunctive score is the coverage-weighted geometric mean

$$S_{\text{conj}} = \exp\left(\frac{\sum_{v \in V} w_v \ln u_v}{\sum_{v \in V} w_v}\right) \times \underbrace{\frac{|V|}{|V_{\text{all}}|}}_{\text{coverage}},$$

where $w_v$ are per-view weights (evolutionary views (episodic selection, presence/absence homoplasy and reconciliation-inferred transfer) are weighted 1.25; mobility and protein-language-model novelty 1.0) and the coverage factor down-weights families supported by few views. Because the geometric mean is small unless *every* contributing view is high, the rule is conjunctive: a family must be flagged by several orthogonal axes to score near the top.

#### Simulation benchmark

The controlled benchmark (`test/00_simulate.py`, fixed seed) generates a pangenome of 300 gene families with 20 planted determinants and decoy families constructed so that each decoy trips exactly one view (e.g. a horizontally transferred but non-resistance gene, or a positively selected core gene). This design deliberately stresses the conjunction: no single view separates determinants from decoys, so recovery is attributable to the multi-view combination rather than to any one signal.

#### Notes and extensions

**Per-view feature definitions.** Each view reduces a gene family (type-A) or a core-genome site (type-B) to a single rank-normalized score.

*Protein-language-model novelty:* ESM-2 (`esm2_t33_650M_UR50D`) embeddings of the family representatives, mean-pooled over residues, whitened by PCA (to decorrelate inputs), and scored as the mean of three rank-normalized unsupervised anomaly estimators: mean distance to the $k$=15 nearest neighbors, an Isolation Forest anomaly score, and squared distance to the whitened centroid.

*Mobility/composition:* for each member, the deviation of GC content and of codon-usage frequencies from that member's own host genome, averaged over the family.

*Episodic selection:* a HyPhy BUSTED gene-wide test acts as a gate, after which MEME, FEL, FUBAR (sites) and aBSREL (branches) run on the families that pass; the summary score is dominated by the aBSREL selected-branch fraction.

*Presence/absence homoplasy:* the Fitch-parsimony step count of the binary presence/absence character on the species tree (excess steps above one count independent gains/losses), restricted to accessory families.

*HGT reconciliation:* AleRax gene-tree/species-tree reconciliation under the UndatedDTL model (which specifically accounts for Duplication, Transfer (horizontal gene transfer), and Loss (DTL) events in an undated framework.), summarized by the per-family inferred transfer count and rate.

The five scores are combined as described in *Conjunctive fusion score.*

**Toward a learned combiner.** The fixed, hand-specified weights are the binding limitation on real data: the ablation (Table S1; Fig 2F) shows that the single presence/absence-homoplasy view is more predictive than the equal-weighted conjunction, and that including the anti-correlated episodic-selection view lowers AUPRC. The natural extension, outlined in the Discussion, would be to replace the fixed geometric mean with a combiner trained on a subset of labeled determinants (for example logistic regression or a learning-to-rank model over the five per-view scores). This would then be evaluated with leave-class-out cross-validation, so that uninformative or anti-correlated views are down-weighted automatically rather than by hand. This is outside the scope of this work, and is left to future work.

**Robustness and sensitivity.** Two diagnostics bound the principal failure modes. The negative-control simulation (Fig S9A) confirms that neutrally evolved pangenomes do not generate spurious presence/absence convergence, so the enrichment seen on real data is not an artefact of the scoring. The sampling-sensitivity analysis (Fig S9B) quantifies how the convergence view depends on sampling depth: subsampling the real pangenomes and recomputing the presence/absence homoplasy score raises AUROC from 0.60 to 0.66 (*E. coli*) and from 0.67 to 0.71 (*K. pneumoniae*) between 25 and 150 genomes, with the curve flat beyond ~75 genomes. Sampling depth is therefore not the factor limiting recovery at $n = 150$; notably, the convergence view carries signal in *both* species, even where the equally weighted conjunction does not.

**Table S1: Full ablation ladder** (simulated benchmark): AUPRC, AUROC, $\mathrm{BEDROC}_{20}$, top-1% enrichment factor (EF), and recall in the top 200 for every view subset and fusion rule evaluated.

| Configuration | Fusion | AUPRC | AUROC | $\mathrm{BEDROC}_{20}$ | $\mathrm{EF}_{1\%}$ | $R \in 200$ |
|---|---|---|---|---|---|---|
| novelty + conv_pa | conjunctive | 1.00 | 1.00 | 1.00 | 15.0 | 1.00 |
| full (5-view) | conjunctive | 1.00 | 1.00 | 1.00 | 15.0 | 1.00 |
| drop novelty | conjunctive | 1.00 | 1.00 | 1.00 | 15.0 | 1.00 |
| drop mobility | conjunctive | 1.00 | 1.00 | 1.00 | 15.0 | 1.00 |
| drop conv_pa | conjunctive | 1.00 | 1.00 | 1.00 | 15.0 | 1.00 |
| drop hgt | conjunctive | 1.00 | 1.00 | 1.00 | 15.0 | 1.00 |
| drop selection | conjunctive | 1.00 | 1.00 | 1.00 | 15.0 | 1.00 |
| novelty + hgt | conjunctive | 1.00 | 1.00 | 1.00 | 15.0 | 1.00 |
| full (5-view) | additive | 1.00 | 1.00 | 1.00 | 15.0 | 1.00 |
| novelty + mobility | conjunctive | 0.99 | 1.00 | 0.99 | 15.0 | 1.00 |
| novelty + selection | conjunctive | 0.98 | 1.00 | 0.98 | 15.0 | 1.00 |
| selection | additive | 0.75 | 0.95 | 0.80 | 15.0 | 1.00 |
| mobility | additive | 0.72 | 0.98 | 0.82 | 10.0 | 1.00 |
| conv_pa | additive | 0.67 | 0.98 | 0.76 | 10.0 | 1.00 |
| hgt | additive | 0.67 | 0.98 | 0.78 | 10.0 | 1.00 |
| novelty | additive | 0.66 | 0.98 | 0.73 | 10.0 | 1.00 |

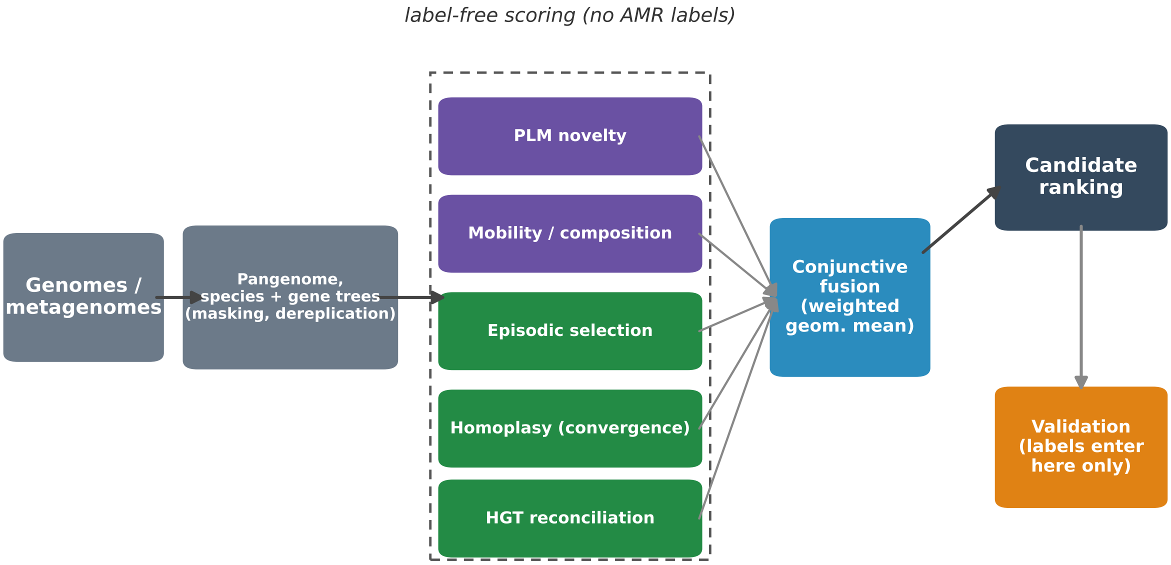


**Figure S1: Overview of the label-free pipeline.** Genomes are reduced to a pangenome with a species tree and per-family gene trees. Five label-free views (protein-language-model novelty, mobility/composition, episodic selection, homoplasy, and reconciliation-inferred HGT) are scored without any resistance label and combined by a conjunctive fusion. AMR labels enter only at the validation step.

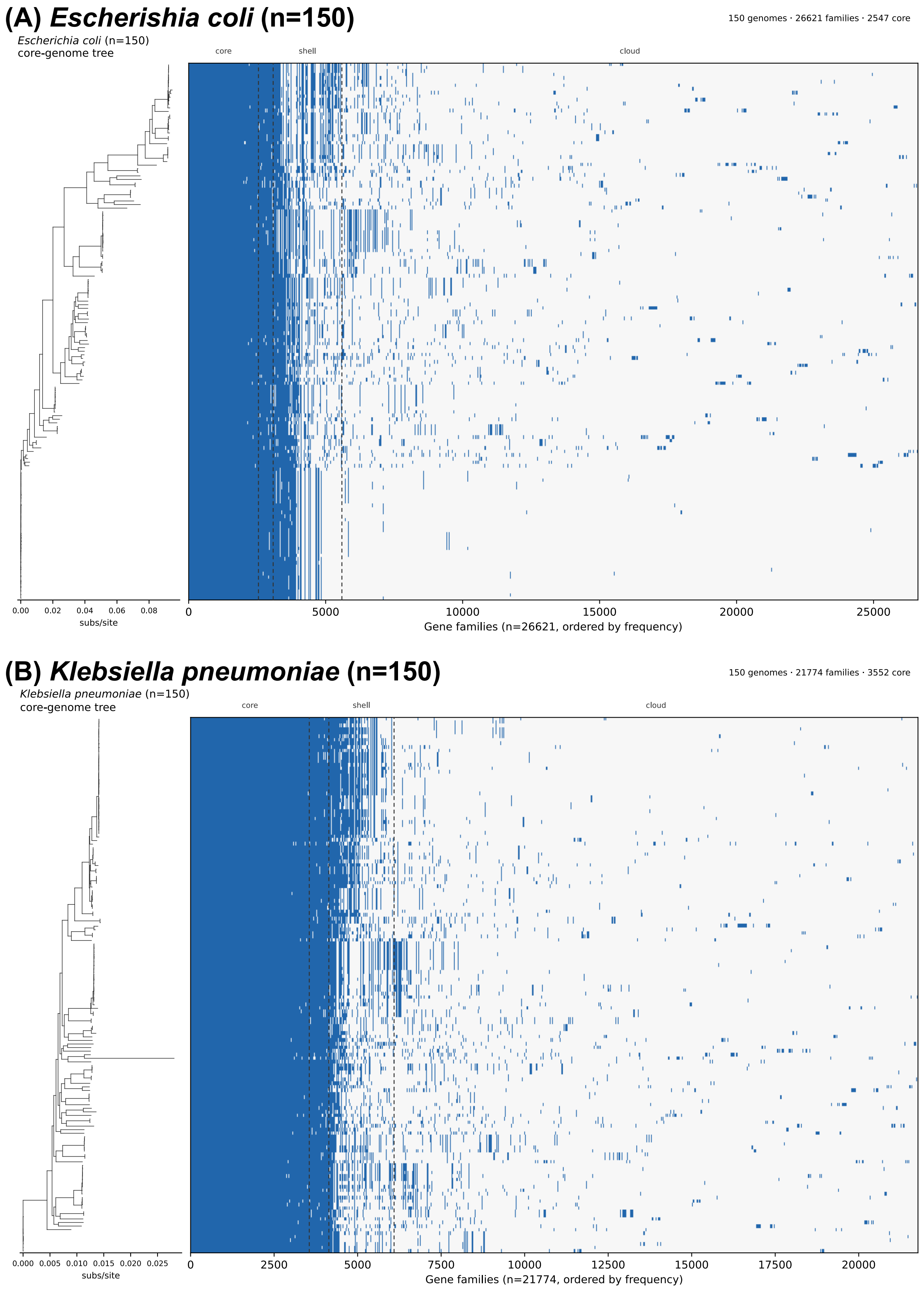


**Figure S2: Core-genome phylogeny and gene presence/absence.** For (A) *E. coli* and (B) *K. pneumoniae*, the maximum-likelihood core-genome tree (left) is aligned row-for-row with the binary gene presence/absence matrix (right; genomes in tree order, gene families ordered by frequency). The solid left block is the core genome; the accessory compartment to the right is structured by clade. Dashed vertical lines mark the core/soft-core/shell/cloud partition boundaries.

#### (A) *Escherishia coli*

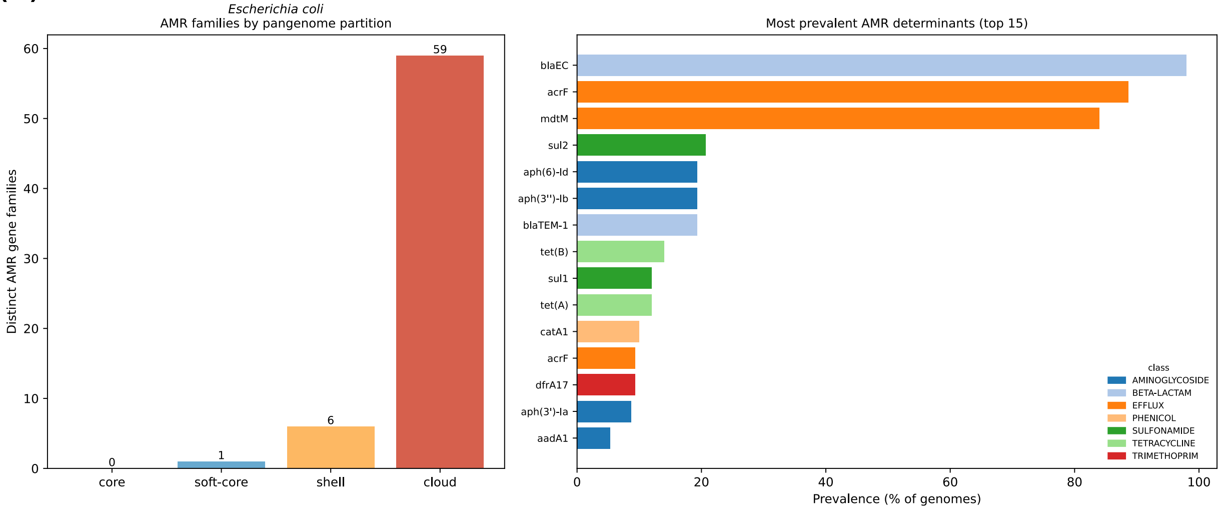


#### (B) *Klebsiella pneumoniae*

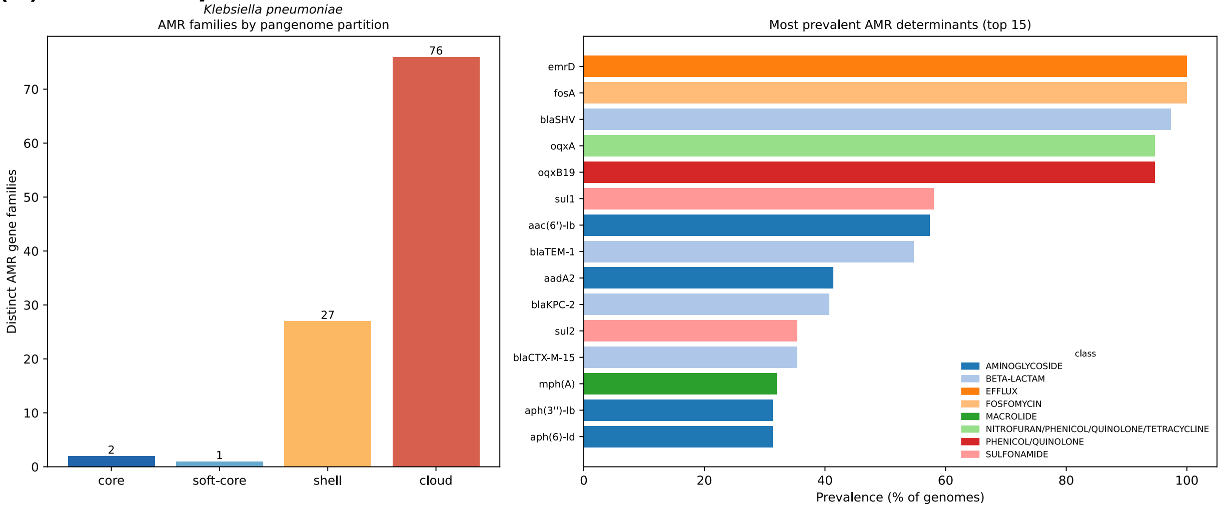


**Figure S3: Known resistance determinants are concentrated in the accessory pangenome.** AMRFinderPlus hits on the gene-family representatives. *Left:* number of distinct determinant families per pangenome partition (almost all in shell/cloud). *Right:* most prevalent determinants (% of genomes), coloured by resistance class. Core/soft-core hits are intrinsic genes (e.g. $bla_{\mathrm{EC}}$ in *E. coli*; *fosA*, $bla_{\mathrm{SHV}}$ in *K. pneumoniae*).

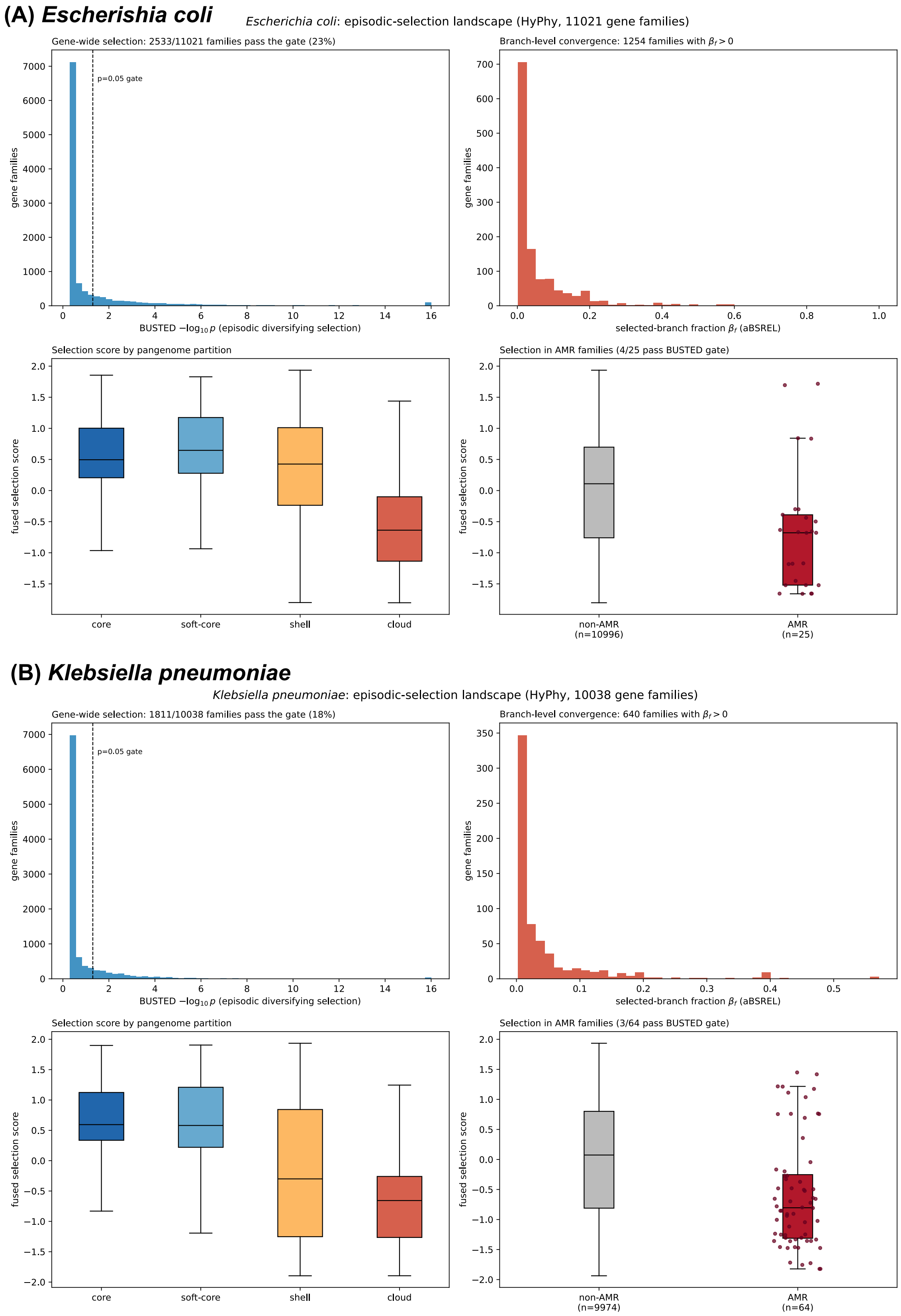


**Figure S4: Episodic-selection landscape of the pangenome** (HyPhy), for (A) *E. coli* (11,021 families) and (B) *K. pneumoniae* (10,038 families). Within each panel, from top left: BUSTED gene-wide significance (23% and 18% of families pass the $p < 0.05$ gate, respectively); aBSREL selected-branch fraction; fused selection score by pangenome partition; and fused selection score in AMR vs non-AMR families, where known determinants score below the genome background in both species.

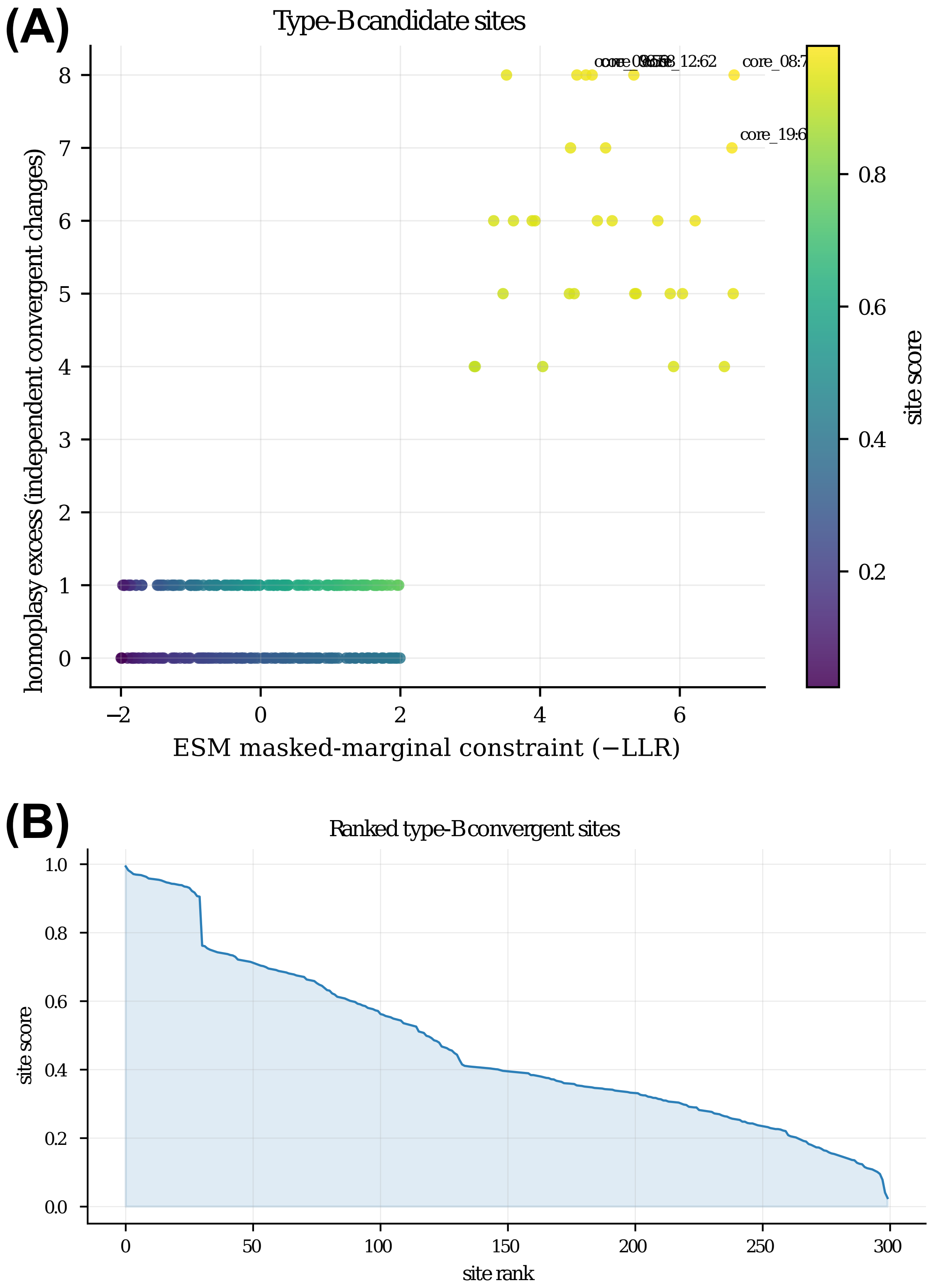


**Figure S5: Type-B (point-mutation) track.** (A) Each core-gene site by ESM-2 masked-marginal constraint and homoplasy excess, colored by combined site score. (B) Ranked site scores. Note how the 20 planted convergent sites (benchmark) separate from the background.

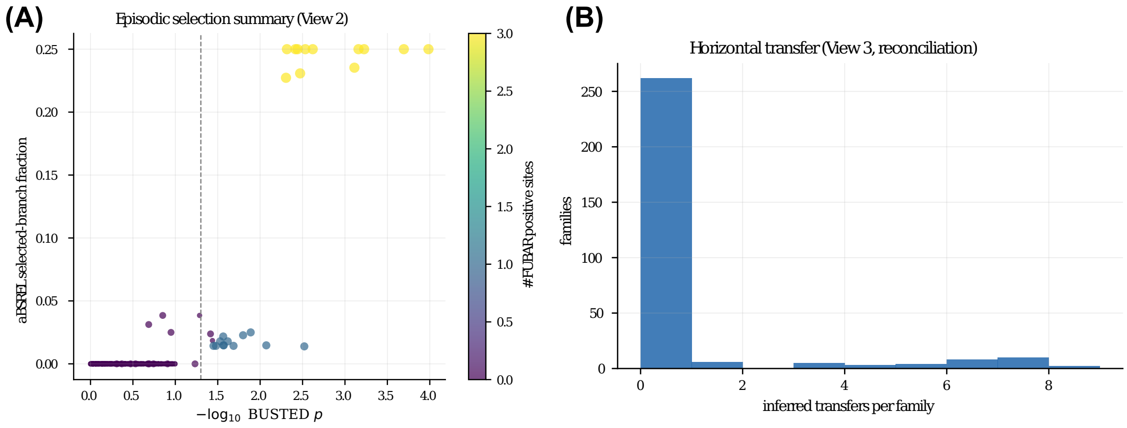


**Figure S6: Evolutionary view diagnostics.** (A) Episodic-selection summary: aBSREL selected-branch fraction vs BUSTED gene-wide significance. (B) Distribution of inferred horizontal transfers per family.

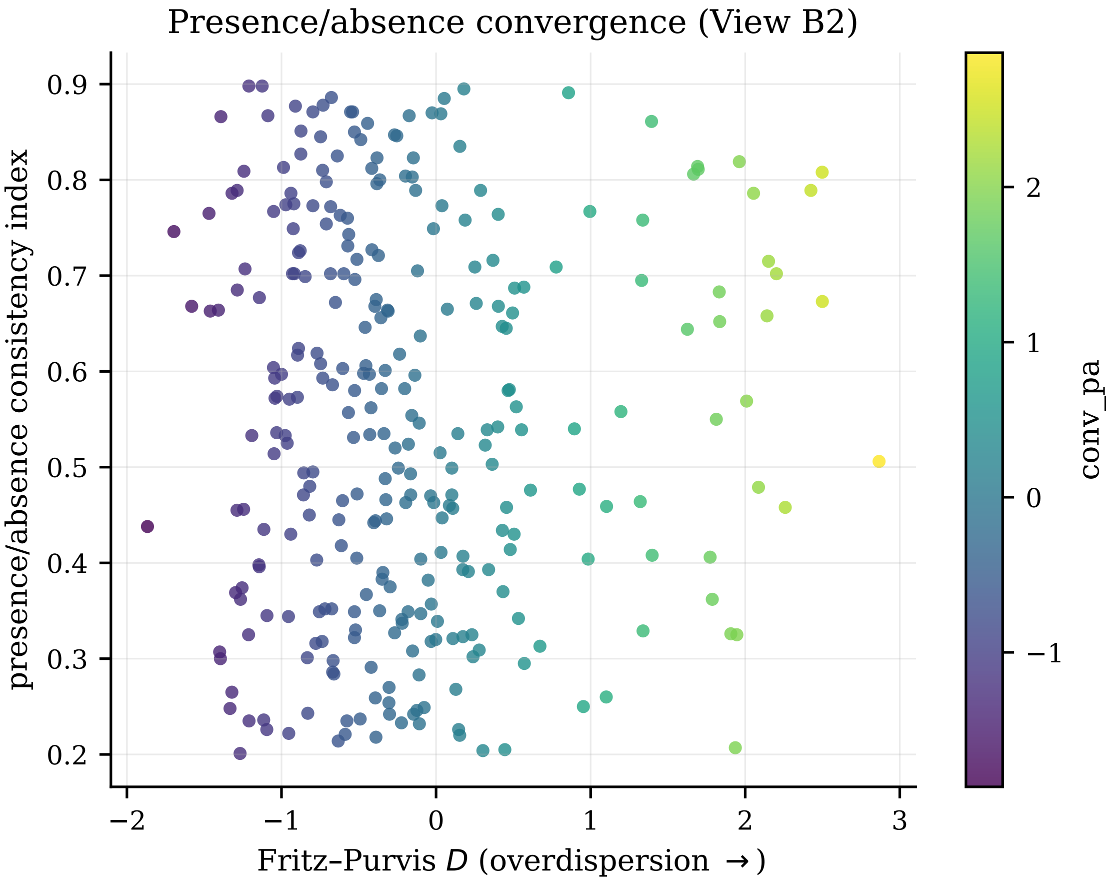


**Figure S7: Presence/absence convergence.** Presence/absence consistency index per accessory family vs Fritz-Purvis $D$, colored by the convergence score. Overdispersed (HGT-like) families lie to the right.

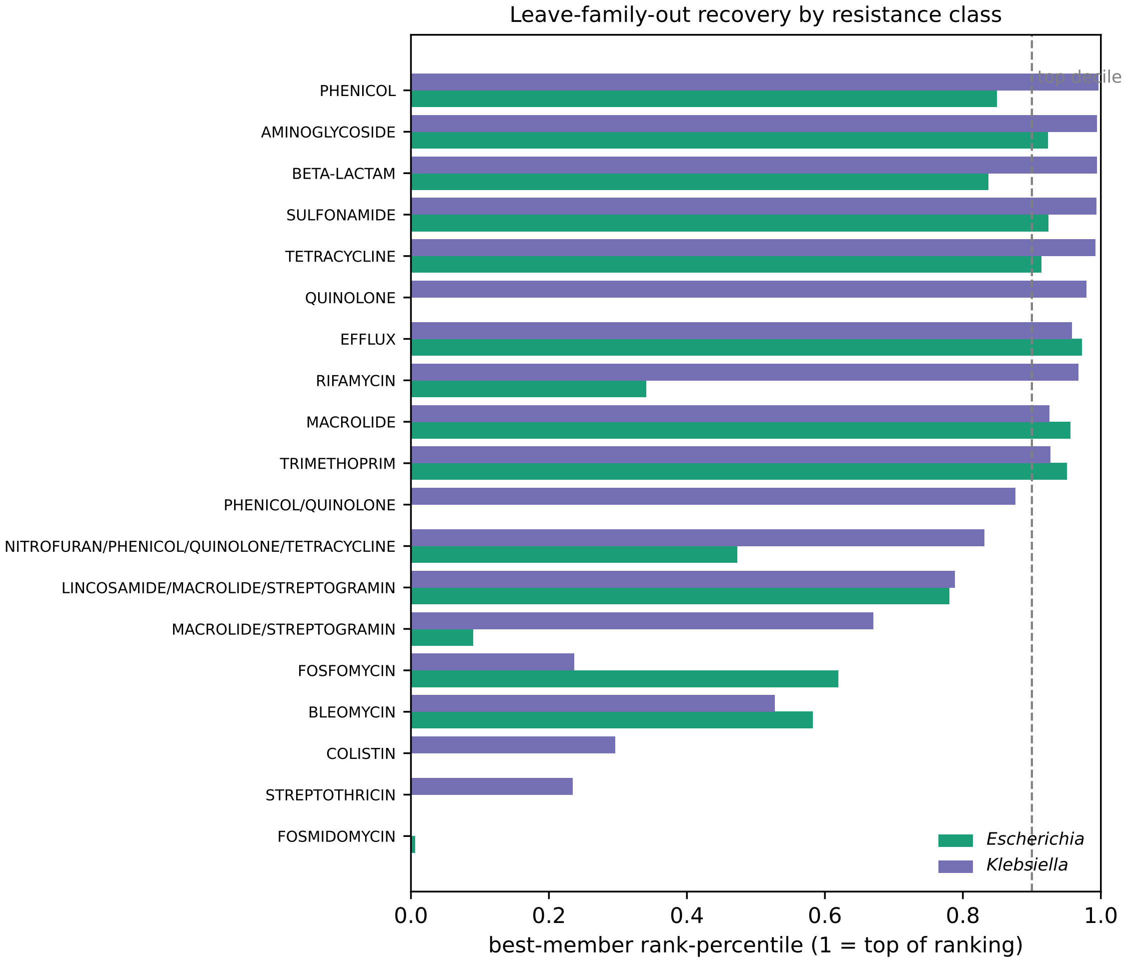


**Figure S8: Leave-family-out recovery by resistance class** (real data). For each AMRFinderPlus resistance class, the rank-percentile of the best-scoring family member under the label-free conjunction (1 = top of the ranking; dashed line, top decile), for *E. coli* and *K. pneumoniae*. The best member of most classes reaches the top decile in *K. pneumoniae* (10 of 18 classes), but fewer in *E. coli* (6 of 15).

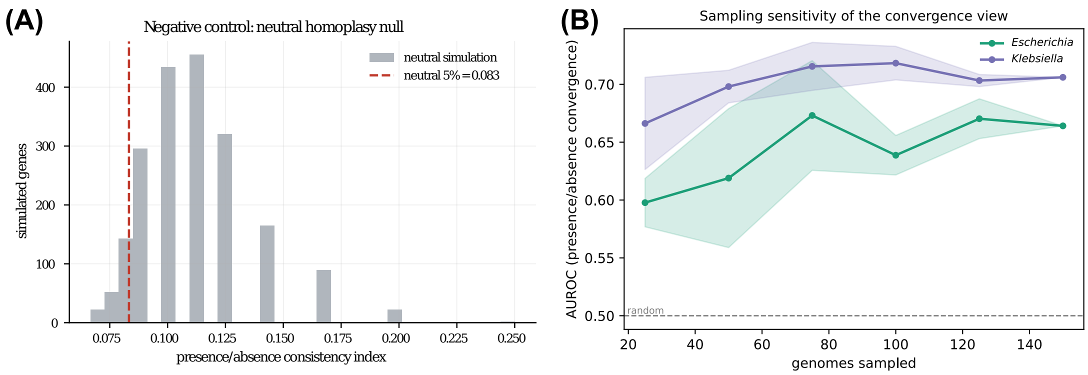


**Figure S9: Negative control and sampling sensitivity.** (A) Neutral-null distribution of the presence/absence consistency index; true determinants are expected below the 5th-percentile line. (B) Sampling sensitivity on real data: AUROC of the presence/absence-convergence view against AMRFinderPlus labels as a function of the number of genomes subsampled (mean $\pm$ s.d. over five random replicates per size; accessory families only, single view). Recovery rises steeply up to $\sim$75 genomes and then plateaus in both species, indicating that the 150-genome sampling depth used here is not limiting for this view.
